# Vulnerable species interactions are important for the stability of mutualistic networks

**DOI:** 10.1101/604868

**Authors:** Benno I. Simmons, Hannah S. Wauchope, Tatsuya Amano, Lynn V. Dicks, William J. Sutherland, Vasilis Dakos

## Abstract

Species are central to ecology and conservation. However, it is the interactions between species that generate the functions on which ecosystems and humans depend. Despite the importance of interactions, we lack an understanding of the risk that their loss poses to ecological communities. Here, we quantify risk as a function of the vulnerability (likelihood of loss) and importance (contribution to network stability in terms of species coexistence) of 4330 mutualistic interactions from 41 empirical pollination and seed dispersal networks across six continents. Remarkably, we find that more vulnerable interactions are also more important: the interactions that contribute most to network stability are those that are most likely to be lost. Furthermore, most interactions tend to have more similar vulnerability and importance across networks than expected by chance, suggesting that vulnerability and importance may be intrinsic properties of interactions, rather than only a function of ecological context. These results provide a starting point for prioritising interactions for conservation in species interaction networks and, in areas lacking network data, could allow interaction properties to be inferred from taxonomy alone.

## Introduction

Species are the predominant biological unit of interest across ecology and conservation. However, it is interactions between species, rather than species themselves, that mediate the ecological functions that drive community dynamics and support biodiversity (1). For example, pollination interactions shape co-evolution in diverse plant-animal communities (2), while seed dispersal maintains spatial patterns of diversity (3). Given the importance of interactions for ecosystem functioning, their loss could have reverberating effects on entire communities and, ultimately, the ecosystem services they deliver (4, 5).

Interactions are thus a vital component of biodiversity, but they remain largely neglected (6). Studies tend to focus on the impact of anthropogenic stressors on single interactions at single sites (7), while the few studies that have considered interaction loss at the community level are either at local scales (8), based on hypothetical network structures (9) or only consider aggregate properties of interactions, rather than considering them individually (10). There is thus an urgent need to incorporate interactions into studies assessing community responses to environmental change. Specifically, we lack a quantitative understanding of the risk that interaction loss poses to communities, which, in turn, limits our ability to make conservation decisions.

Here, we address this gap by quantifying the risk of interaction loss to 41 pollination and seed dispersal communities that, combined, comprise a global dataset of 4330 species-species links (see Materials and Methods). Such mutualisms are fundamental to the functioning of most communities. The loss of pollination can lead to pollen limitation, potentially compromising reproduction for the vast majority of plant species that rely, to some extent, on animal pollinators (11, 12). Similarly, the disruption of seed dispersal can have deleterious, cascading consequences for those woody plant species that depend on frugivores, which can exceed 90% in biodiverse ecosystems such as tropical rainforests (13).

Conventionally, risk is a function of both the likelihood of an event occurring and the severity of the impacts if it did occur (14). For ecological networks, we therefore start by reasoning that the risk of losing a particular link is a function of (i) the likelihood of that link being lost (link vulnerability, *V*), and (ii) the severity of the consequences to the community if the link is lost (link importance, *I*). Using novel quantitative methods, we calculate the vulnerability and importance of all links in our dataset (Figure 1) and show that the most vulnerable links in a community are also those that contribute most to its structural stability (Materials and Methods). We next examine whether vulnerability and importance are intrinsic attributes of interactions, rather than functions of ecological context, by testing whether an interaction’s vulnerability and importance is more similar across occurrences than expected by chance. Our aim is to explore the risk of interaction loss in mutualistic communities and to inform their conservation. Hereafter we distinguish between the terms *interaction* and *link*: *interaction* refers to all occurrences of a given taxon-taxon interaction identity, while *link* refers to a single occurrence of an *interaction* in a particular network. Thus, for example, the *Bombus pratorum* – *Leucanthemum vulgare interaction* is present in our data, while *Bombus pratorum* – *Leucanthemum vulgare links* occur in two networks.

**Figure 1:**
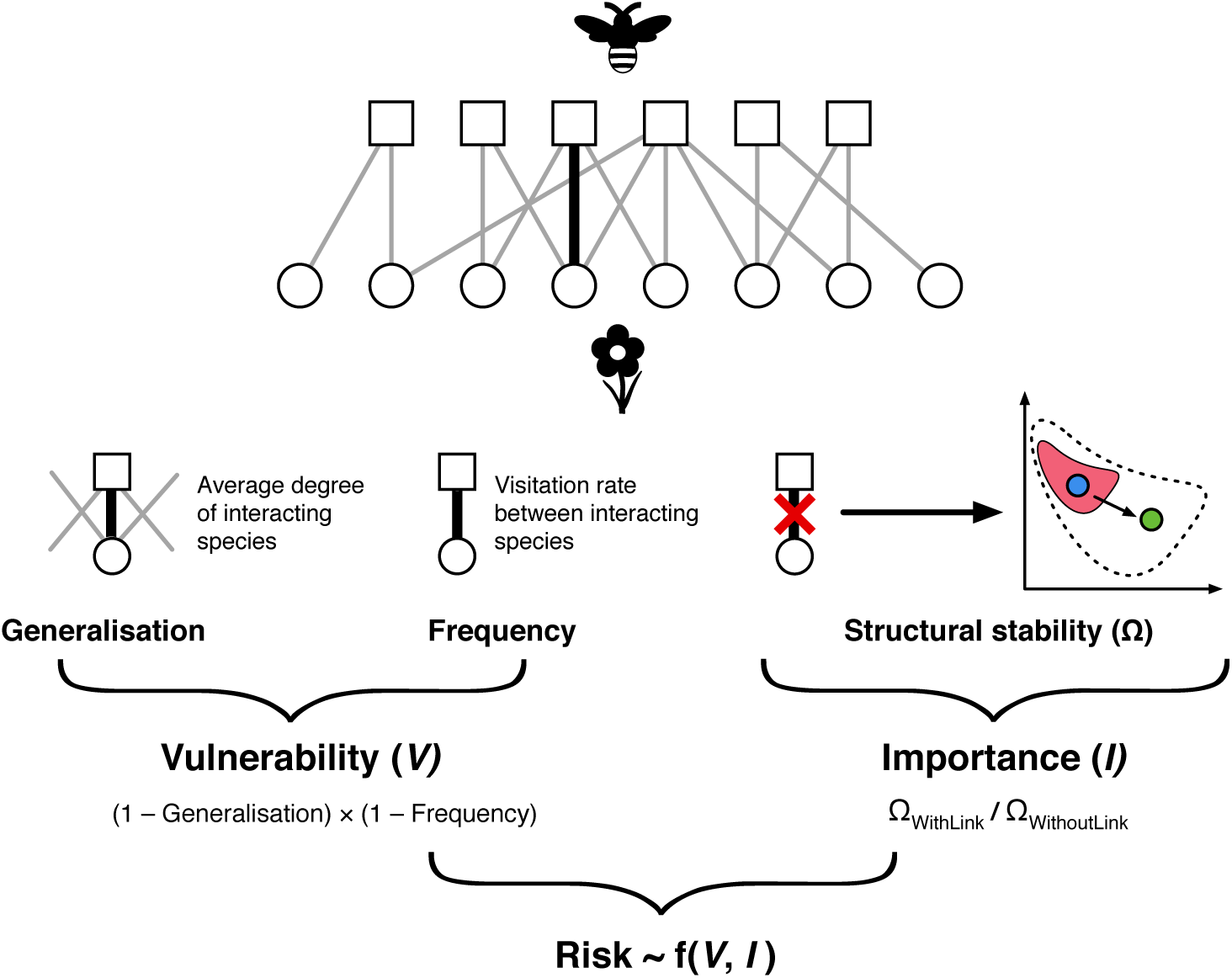
The quantities used in the analysis. Top: an illustrative network depicting interactions between plants and pollinators. A focal link is highlighted in black. The generalisation of a link is the average degree of the interacting species. The frequency of the link is the visitation rate between the interacting species. Combined, these measures determine the vulnerability of a link, such that vulnerable links are low frequency interactions between specialists. The structural stability of the network was measured with the focal link and without the focal link. The ratio of these two values is the importance of the link. This is represented by the graph on the right of the figure. The structural stability of a network is defined as the parameter space of intrinsic growth rates in which all species in a community can have positive abundances. This is represented for the whole community by the dotted outline and for the whole community without the focal link by the pink shape. In this case, removing the link has reduced the structural stability of the community (reduced the size of the shape). This means that a perturbation that moves the community from the initial state (blue circle) to a final state (green circle) will result in extinctions without the focal link because the community moves outside the pink feasibility domain. However, no extinctions would occur under the same perturbation in the original community before the focal link was removed, because the final state of the community (green circle) is within the original feasibility domain (dotted outline). Together, vulnerability and importance describe the risk to a community of losing a particular link.

## Results

### Relationship between link vulnerability and importance

We calculated the vulnerability and importance of 4330 links from a global dataset of 29 plant-pollinator and 12 plant-seed disperser networks (Figure 1). Vulnerability was measured as a function of link frequency (how often the two species involved in the link interact) and link generalisation (the mean number of links [the mean degree] of the two species involved in the link) (8). This means that weak (less frequent) links between specialists were more vulnerable than strong (more frequent) links between generalists (8). Importance was defined as the contribution of a given link to the feasibility of a network, where feasibility is a measure of a network’s ability to withstand environmental variation without leading to species extinctions (15, 16). Important links were those that, when removed, lowered a network’s feasibility; that is, when removed, they reduced the amount of environmental variability a network can tolerate before species extinctions took place (Figure 1; see Materials and Methods). We found that there was a significantly positive correlation between the vulnerability and importance of links across the 41 networks (Wald test: χ^2^ = 53.71, df = 1, P = < 0.001) (Figure 2). This correlation indicates that the links that contribute most to the structural stability of communities are those links that are most likely to be lost in the face of environmental changes.

**Figure 2:**
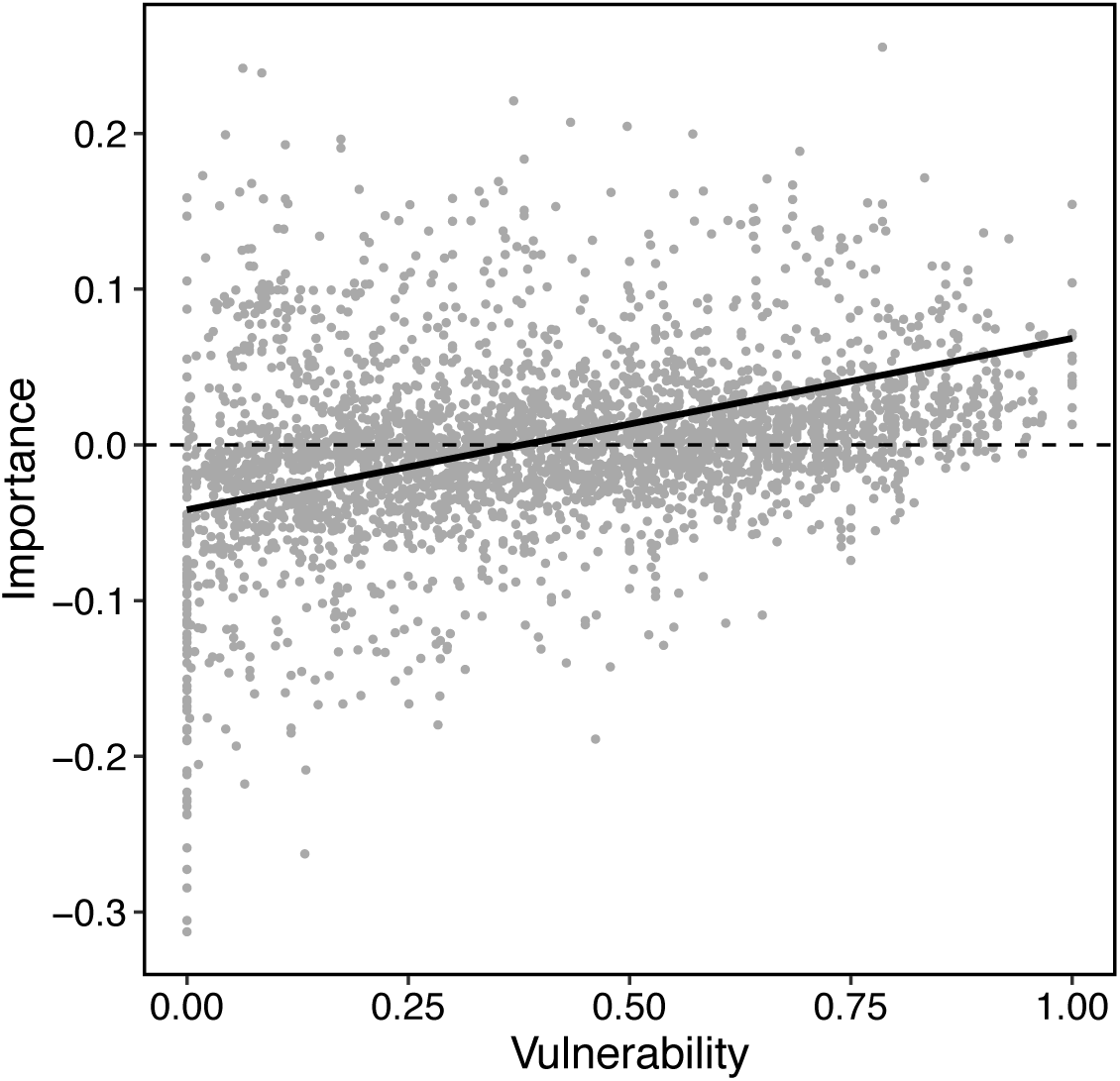
The relationship between vulnerability (the likelihood of a link being lost) and importance (the contribution of a link to a network’s structural stability) for all 3391 species-species links across 41 mutualistic networks. Best fit line is from a mixed effects model with importance as the response variable, vulnerability as a fixed effect, and network identity as a random effect.

### Taxonomic consistency of link vulnerability and importance

A consistent, positive relationship between vulnerability and importance could arise if links tend to have the same vulnerability and importance independent from the network in which they occur. This would imply some form of evolutionary conservatism in interaction properties. We tested this hypothesis by assessing the extent to which vulnerability and importance exhibited taxonomic consistency: the tendency for an interaction’s vulnerability and importance to be more similar across all the networks in which the interaction occurs than expected by chance. If vulnerability and importance exhibit taxonomic consistency, then all occurrences of a given interaction should have similar levels of these properties. For each interaction, we compared the variance in vulnerability and importance to a null expectation where links were sampled randomly (see Materials and Methods). We carried out analyses at genus, family and order levels, but not at the species level, because very few interactions at the species level occurred more than once in the data. We found a strong tendency towards consistency for both vulnerability and importance at all taxonomic levels (between 76% and 83% of interactions had more similar values of vulnerability and importance than expected by chance; Figure 3). Considering vulnerability, there was significant taxonomic consistency for 18% of genus, 17% of family and 30% of order interactions (see Materials and Methods). For importance, interactions had significant taxonomic consistency for 14% of genus, 20% of family and 33% of order level interactions. Conservatism was observed across large geographic scales, with many significantly-consistent interactions comprised of links occurring in different regions or continents. These results suggest that vulnerability and importance may be, to some extent, intrinsic properties of interactions and not only a function of ecological context.

**Figure 3:**
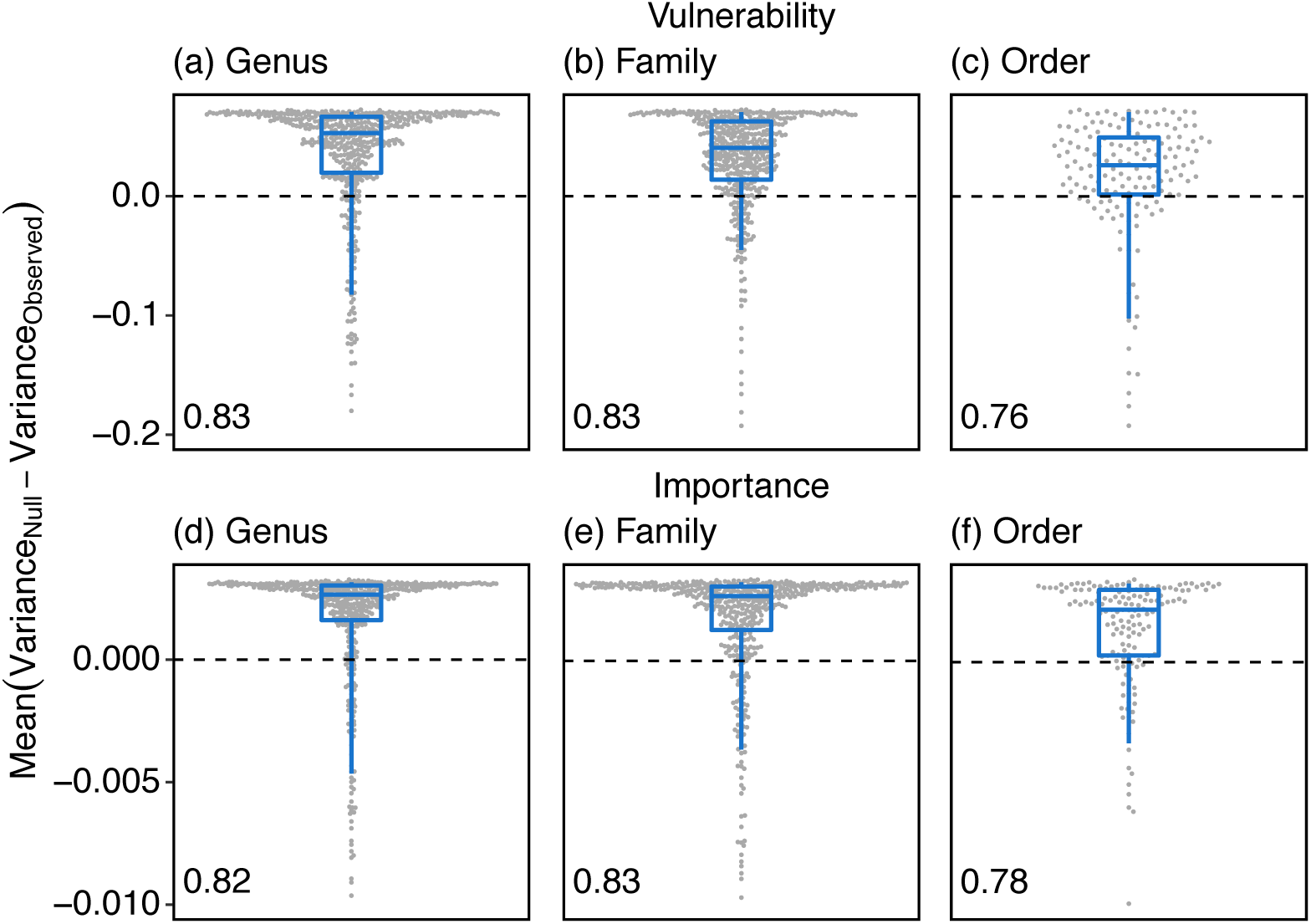
The degree of taxonomic consistency for each interaction at genus (*n* = 469), family (*n* = 466) and order (*n* = 151) levels, for both vulnerability (likelihood of a link being lost) and importance (contribution of a link to a network’s structural stability). Taxonomic consistency is the tendency for properties of an interaction to be more similar across occurrences than expected by chance. Points represent individual interactions. Boxplots represent 5%, 25%, 50%, 75% and 95% quantiles of the same data, moving from the bottom whisker to the top whisker. Number in bottom left of each panel is the proportion of interactions which exhibited positive consistency (Variance_Observed_ < Variance_Null_). For visualisation, a small number of points with low values were removed. The percentage of points with values lower than the y-axis minimum are as follows for each panel: (a) 1.5%, (b) 1.1%, (d) 3.2%, (e) 1.5%, (f) 1.3%.

### Mapping the risk of link loss

The two components of risk – vulnerability and importance – reflect, respectively, how likely a link is to be lost and how serious the consequences of that loss are for the community. Using an illustrative plant-seed disperser network, we coloured links based on their vulnerability and importance, and highlighted those that exhibited taxonomic consistency (Figure 4). As expected, given the positive correlation between vulnerability and importance, we find that a substantial proportion of links are either highly vulnerable and contribute strongly to structural stability (22.5%; dark red in Figure 4) or have low vulnerability and contribute negatively to stability (19.4%; light purple in Figure 4). Conversely, few links are of low vulnerability and positive importance (5.4%; light yellow in Figure 4) or high vulnerability and negative importance (2.3%; dark purple in Figure 4). Identifying those links that are vulnerable and which benefit community stability as a whole provides a potential starting point when deciding which links should be priorities for conservation (Figure 4). If two links have similar vulnerability and importance, but one exhibits stronger taxonomic consistency than the other, then the more consistent link may be of higher concern as it could be important and susceptible to extinction across communities in different geographical regions.

**Figure 4:**
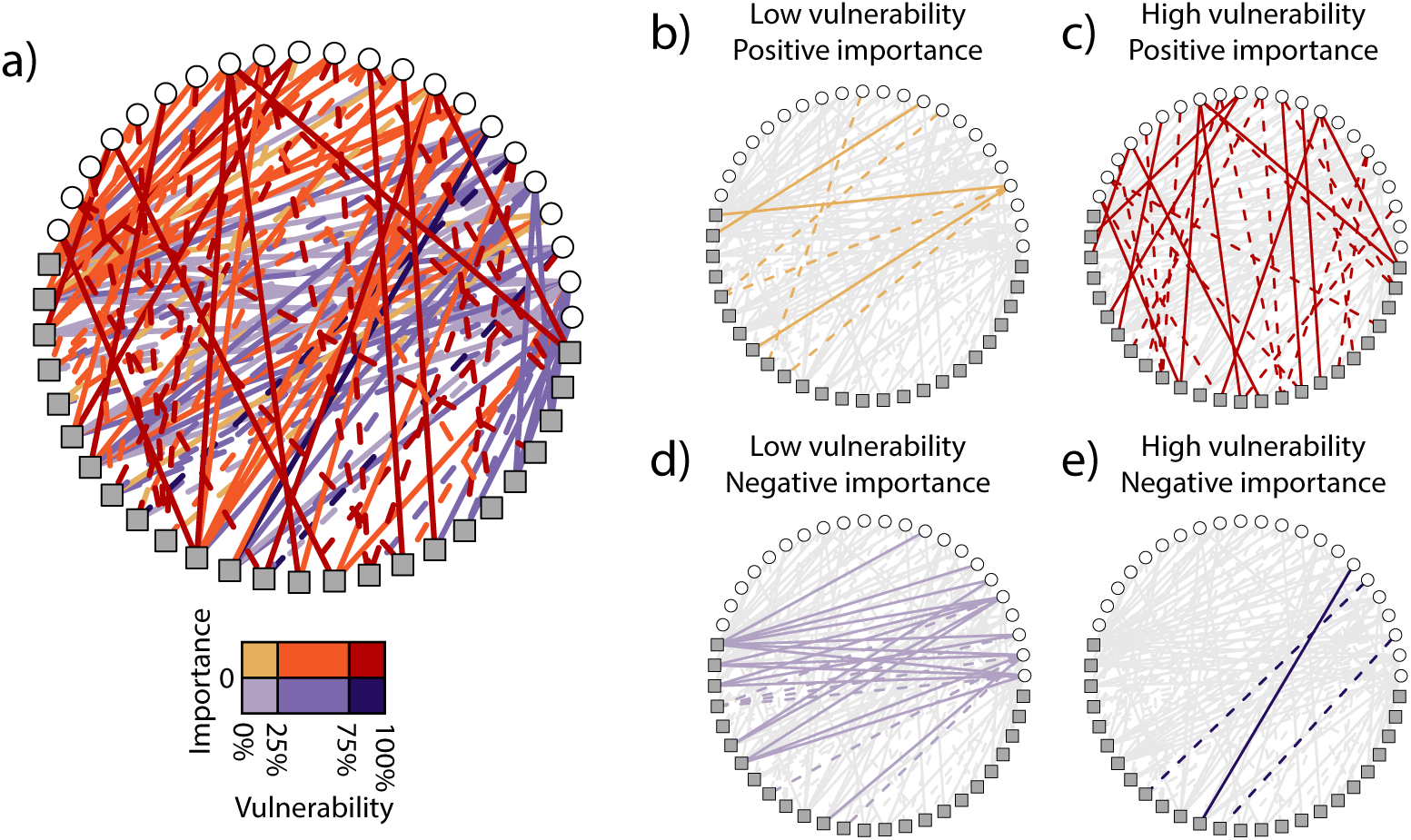
Mapping vulnerability, importance and conservatism onto an example plant-seed disperser network (squares are plant species; circles are seed disperser species). Solid lines indicate significant taxonomic consistency at the genus, family or order level. Links are categorised based on (i) whether they positively or negatively contribute to the feasibility of the community; that is whether their importance (contribution to network feasibility) is positive or negative, and (ii) whether their vulnerability is in the bottom 25% of vulnerability values, the middle 50% or the top 25%. (a) Shows all links together; (b) highlights low vulnerability, positive importance links; c) highlights high vulnerability, positive importance links; (d) highlights low vulnerability, negative importance links; (e) highlights high vulnerability, negative importance links. We suggest that consistently-important and vulnerable links (solid, dark red lines) should be high priorities in conservation.

## Discussion

Our analysis represents the first attempt to quantify the risk that link loss poses to ecological communities. We find that, across 41 ecological networks, the links that contribute most to a network’s ability to tolerate environmental perturbations are the same links that are most likely to be lost in the face of such perturbations (Figure 2). We additionally find that there is a strong tendency for interactions to have more similar vulnerability and importance across occurrences than expected by chance, with a substantial proportion of interactions exhibiting this signal significantly (Figure 3). By combining these results, we are able to map the risk of interaction loss onto empirical communities, which could be used to guide conservation efforts (Figure 4).

The positive relationship between vulnerability and importance means that the more vulnerable a link is, the more likely it is to have a negative impact on network feasibility if it is lost. Vulnerability is therefore an important indicator of the extent to which a link supports or hinders a community’s ability to tolerate environment variation and, thus, species’ long-term persistence. From a conservation perspective, this result is concerning as it suggests that losing vulnerable interactions reduces the ability of mutualistic networks to absorb future stressors. However, it also suggests that our proposed link vulnerability measure enables estimates of how much a link benefits a community (which may be difficult to measure otherwise) using only simple topological information. Aizen et al (8) found that vulnerable links were found in places with greater habitat loss. Combined with our results, this suggests environmental stressors like habitat loss may be detrimental for whole-community stability, not just those links which are vulnerable.

That those links that are stronger contributors to structural stability are more vulnerable to extinction mirrors findings that *nodes* that contribute most to a network’s nestedness (and thus persistence) are those that are most likely to go extinct (17). The positive relationship between link vulnerability and importance suggests that links have a tendency to fall into one of two categories: low vulnerability and low importance or high vulnerability and high importance. Thus, some links have a high probability of survival to the detriment of the network as a whole, while others contribute to the collective good at the expense of their own viability (17, 18). While the causes of these patterns are unclear, perhaps there is a tendency for some species to maximise their fitness by being involved in a mixture of ‘selfless’ links that ensure the community as a whole remains intact, and ‘greedy’ links that provide stable benefit to the species over time. Determining how such ‘selfless links’ arise is an important area for future research as characterising the conditions compatible with their genesis could aid the design or maintenance of resilient ecosystems, and cooperative systems more broadly.

We found that many interactions tend to have similar vulnerability and importance values across occurrences, implying a form of evolutionary conservatism in properties of species interactions. The consistency of vulnerability could be driven by conserved patterns of generalisation or abundance of the interacting partners. For example, pollinator species have been shown to have similar levels of generalisation across their range (19). Similarly, (20) found that particular pollinator clades tended to be generalist, while (21) found significant phylogenetic signal in pollinator interactions. Meanwhile, the taxonomic consistency of importance has substantial implications. While here we consider the importance of individual links, such importance values are governed by the whole network structure, not just the roles of the two partner species involved in the link. To illustrate this, consider two pollinators, *i* and *j*, and two plants, *m* and *n* that all interact with each other; that is *i* and *j* both interact with *m* and *n*. If *j* leaves the network, the importance of all remaining links will change (22). Thus, that link importance is consistent across occurrences suggests that links, and their partner species, are embedded in networks in similar ways, as has been found for species in antagonistic networks (22, 23).

Our results have significant conservation implications. Differences in links’ vulnerability, importance and taxonomic consistency could be used to guide proactive conservation efforts: links with high vulnerability and importance could provide a useful starting point to inform prioritisation before any links are lost (Figure 4). Similarly, highly-important links that are not currently vulnerable could be a focus of monitoring efforts in case they become vulnerable in the future. Importantly, by explicitly focusing on links themselves, our methods may identify high-priority links that are not expected to be so based only on assessments of species extinction risk. Conversely, our results may be able to inform species conservation if high-priority links tend to involve species that are also of high priority. Determining the relationship between the conservation priority of species and the links they form is an important area for future research. Our finding of widespread taxonomic consistency potentially allows properties of interactions to be inferred in regions without network data, even if such properties are only known for congeneric, confamiliar or conorder interactions. This is important because species interaction networks are often cost-and time-intensive to collect, and coverage is highly biased geographically (24, 25).

Conserving links is perhaps even more challenging than conserving species. While species conservation requires one species to remain extant, link conservation requires that two species remain at sufficiently high abundance to still significantly interact: two species must be prevented from going *ecologically* extinct (26–28). Moreover, because interaction extinction often precedes species loss, conservation actions must occur sooner. While link conservation has attracted little attention so far, the importance of interactions like pollination is now widely recognised. Thus, we hope our results can help guide future research in this nascent and important field, because ultimately it is links that support the ecological functions and services that communities provide.

## Materials and Methods

### Data

We assembled a dataset of 4330 plant-animal links from 41 quantitative mutualistic networks spanning a broad geographical range, with data in tropical and non-tropical areas from both islands and mainlands (www.web-of-life.es, [29, 30]). The database spanned two types of mutualism, comprising 3182 pollination links from 29 pollination networks and 1148 seed dispersal links from 12 seed dispersal networks. The data contained 551 plant species and 1151 animal species.

### Interaction properties

#### Link Vulnerability

We developed a measure of network link vulnerability following Aizen *et al*. (8). They identified two factors that determine the vulnerability of a mutualistic link between a plant (*i*) and animal (*j*) species: link frequency (hereafter ‘frequency’) and link degree (hereafter ‘generalisation’) (8). Frequency is how often a link occurs between *i* and *j* (such as the number of times a pollinator species visits a particular flower species), while generalisation is defined as the mean degree of the two species involved in a link, that is, the average number of species with which species *i* and *j* interact (8). This notion of vulnerability aims to capture the sensitivity of a network to the loss of a given link (Figure 1).

We calculated the frequency and generalisation of all links in our dataset. Following Aizen *et al*. (8), we first log_10_ transformed all frequencies. Second, to make results between networks comparable, we standardised frequency and generalisation to between 0 and 1 at the network level. Finally, we calculated the vulnerability of a link between species *i* and *j*, as *V*_*ij*_ = (1 – *f*_*ij*_)(1 – *D*_*ij*_), where *f*_*ij*_ is the standardised link frequency and *D*_*ij*_ is the standardised link generalisation. In this formulation, the index can take values between 0 (least vulnerable) and 1 (most vulnerable), which means that it categorizes weak links between specialist partners as more vulnerable than strong links between generalists.

#### Link Importance

Feasibility is defined as the range of conditions under which all species in a community can stably coexist (16). Feasibility can therefore be thought of as the ‘safe operating space’ of ecological communities: it is an indicator of how much environmental stress a community can tolerate before extinction of any of its constituent species. Formally, feasibility is defined as the volume of the parameter space of intrinsic growth rates in which all species in a community can have positive abundances (31, 32). Feasibility is essential for understanding how communities might respond to future environmental changes. For example, in a very feasible community, there is a large range of conditions under which all species stably coexist. Therefore, in the presence of an environmental perturbation, such as climate change or habitat loss, it is less likely that any of the species in the community decline to extinction. Conversely, in a community with low feasibility, there is a small range of conditions under which all species stably coexist. Therefore, perturbations are more likely to result in species extinctions.

We measured the importance of a link between species *i* and *j* (*I*_*ij*_) as its contribution to the feasibility of a network, defined as the ratio between the feasibility of the network with (*O*) and without (*R*) the focal link: *I*_*ij*_ = Ω _*O*_ / Ω _*R*_, where Ω is the feasibility (Figure 1). Importance values were expressed as ((100 * *I*_*ij*_) – 100), such that *I*_*ij*_ = 0 if the feasibility of the network was identical with and without the focal link. We calculated feasibility following (16). Full details of the mutualistic model and equations used can be found in (16, 32, 33), but we outline these briefly below.

A generalized Lotka-Volterra model of the following form was used:

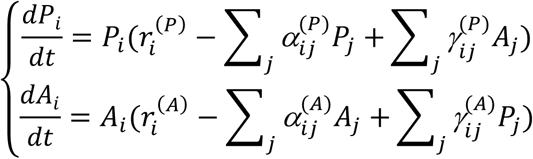

where *P*_*i*_ and *A*_*i*_ give the abundance of plant and animal species *i*, respectively; *r*_*i*_ denotes the intrinsic growth rates; *α* _*ij*_ represents intraguild competition; and *γ*_*ij*_ is the mutualistic benefit. The mutualistic benefit follows the equation 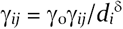, where *γ*_*ij*_ = 1 if there is a link between species *i* and *j* and zero if there is no link; *d*_*i*_ is the degree of species *i*; *δ* is the mutualistic trade-off (34); and *γ*_*o*_ is the overall level of mutualistic strength. A mean field approximation was used for the intraguild competition parameters, setting *α*_*ii*_ ^(*P*)^ and *α*_*ii*_ ^(*A*)^ equal to 1 and a_*ij*_ ^(*P*)^ nd a_*ij*_^(*A*)^ equal to *ρ*(*i ≠ j*). We estimated the mutualistic trade-off, *δ*, empirically across all networks in our dataset. *δ* is given by the slope of two linear regressions (15)

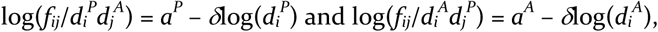

where *f*_*ij*_ is the link frequency between animal species *j* and plant species *i, a*^*P*^ is the intercept for plants and *a*^*A*^ is the intercept for animals. These regressions were performed together on the whole dataset. We obtained a value for *δ* of 0.339, which was consequently used in all simulations. To focus on mutualistic effects, we ran analyses with zero interspecific competition (*ρ* = 0), following (32). Results were qualitatively identical using weak competition (*ρ* = 0.01) (Supplementary Material Figure S1). The average mutualistic strength was set as half the average mutualistic strength at the stability threshold. Contribution to feasibility could not be measured for 931 links which, when removed, resulted in at least one species having no connections; these were excluded from the dataset.

To examine the relationship between interaction vulnerability and importance, we used a linear mixed-effects model, with importance as the response variable, vulnerability as a fixed effect, and network identity as a random effect. Linear mixed-effects models were run and analysed using the ‘lme4’, ‘car’ and ‘MuMIn’ R packages (35–38).

### Taxonomic consistency of vulnerability and importance

We next assessed the extent to which vulnerability and importance exhibited taxonomic consistency: the tendency for an interaction’s vulnerability and importance to be more similar across all the networks in which it occurs than expected by chance. Significant taxonomic consistency would imply a form of evolutionary conservatism: that vulnerability and importance are intrinsic properties of interactions, rather than a function of the ecological context in which they occur (23). We made comparisons at three levels of taxonomic aggregation – genus, family and order – but not at the species level, as only 180 (5.8%) of interactions at the species level occurred more than once in the data (23). For each level of taxonomic aggregation, we excluded all interactions that only occurred once.

If vulnerability and importance exhibit taxonomic consistency, then all occurrences of a given interaction should have similar levels of vulnerability and importance. More specifically, variance in vulnerability and importance across all links of a given interaction should be low. Therefore, for each interaction, we first calculated the variance in vulnerability and importance across all the networks in which the interaction occurred. We then created a corresponding ‘null interaction’, comprising the same number of links as the empirical focal interaction, but consisting of links sampled randomly without replacement from across the dataset. Links that were part of the focal interaction were excluded from this sample. To ensure vulnerability and importance values were comparable between networks, and to control for any network-level effects, we used only relative values of vulnerability and importance. Vulnerability values were rescaled between 0 and 1 at the network level, while importance values were already relative (see definition above). Relative values are more relevant for our study because we were interested in whether interactions tend to have the same relative roles in all communities in which they occur, rather than if they have the same absolute values of a particular property. For example, we wanted to know whether a given link was always the most vulnerable link in a community, rather than if it always has an absolute vulnerability value of, say, 0.7. For each taxon-taxon interaction, at each taxonomic level, we sampled 10,000 null interactions and recorded the mean paired difference between the observed and null variance in vulnerability and importance. If an interaction exhibits taxonomic consistency, the mean paired difference (Variance_Null_ – Variance_Observed_) will be positive, because the observed variance would be lower than that expected by chance. *P* was the probability that a null interaction had lower variance in vulnerability or importance than the observed interaction. Interactions had significant consistency when *P* < 0.05. Taxonomic consistency results were qualitatively identical using weak competition (*ρ* = 0.01) (Supplementary Material Figure S2).

## Author contributions

BIS conceived the idea. BIS and VD designed the analyses. TA assisted with analysis design. BIS conducted all analyses. HSW assisted with analyses. BIS wrote the first draft of the manuscript. All authors discussed the results and contributed to writing the manuscript.

## Acknowledgements

BIS is supported by the Natural Environment Research Council as part of the Cambridge Earth System Science NERC DTP [NE/L002507/1]. HSW was supported by a Cambridge Trust Cambridge-Australia Scholarship and a Cambridge Department of Zoology JS Gardiner Fellowship. TA was supported by the Grantham Foundation for the Protection of the Environment, the Kenneth Miller Trust and the Australian Research Council Future Fellowship (FT180100354). LVD was supported by the Natural Environment Research Council (grants NE/K015419/1 and NE/N014472/1). WJS is funded by Arcadia.

