## Supplementary Information for "Vulnerable species interactions are important for the stability of mutualistic networks"

#### Figure S1: Relationship between link vulnerability and importance ($\rho$ = 0.01)

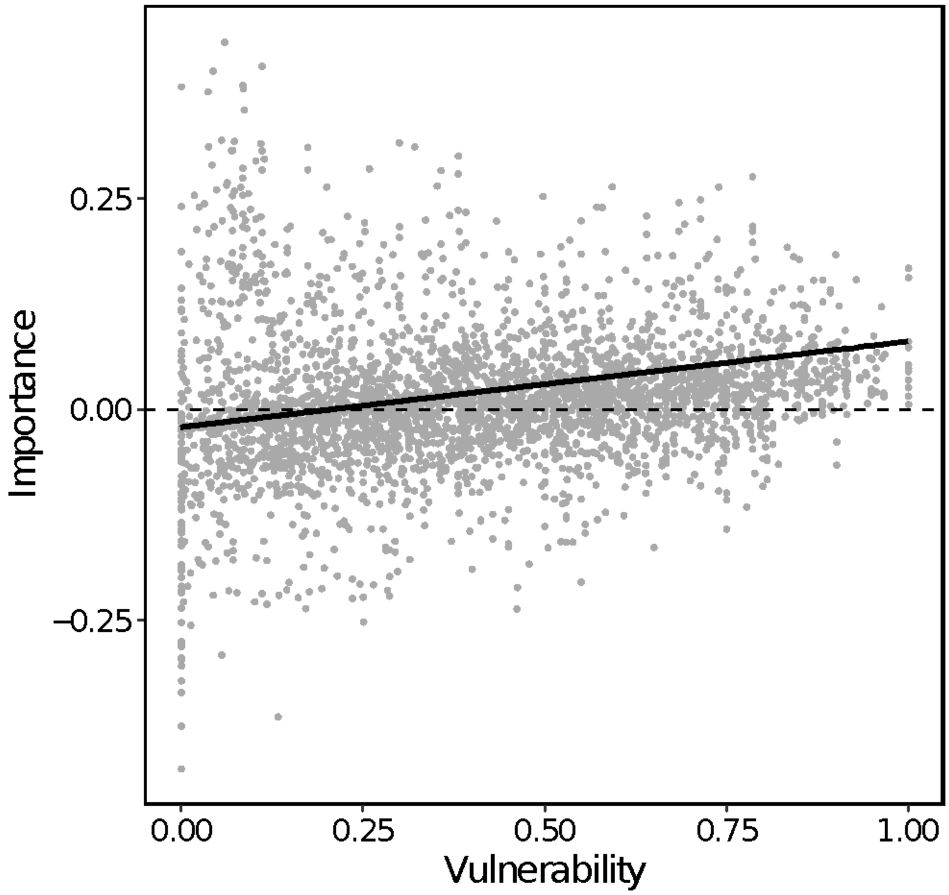


**Figure S1:** The relationship between vulnerability (the likelihood of a link being lost) and importance (the contribution of a link to a network's structural stability) for all species-species links across 41 mutualistic networks. Best fit line is from a mixed effects model with importance as the response variable, vulnerability as a fixed effect, and network identity as a random effect.

### Figure S2: Taxonomic consistency of vulnerability and importance ($\rho$ = 0.01)

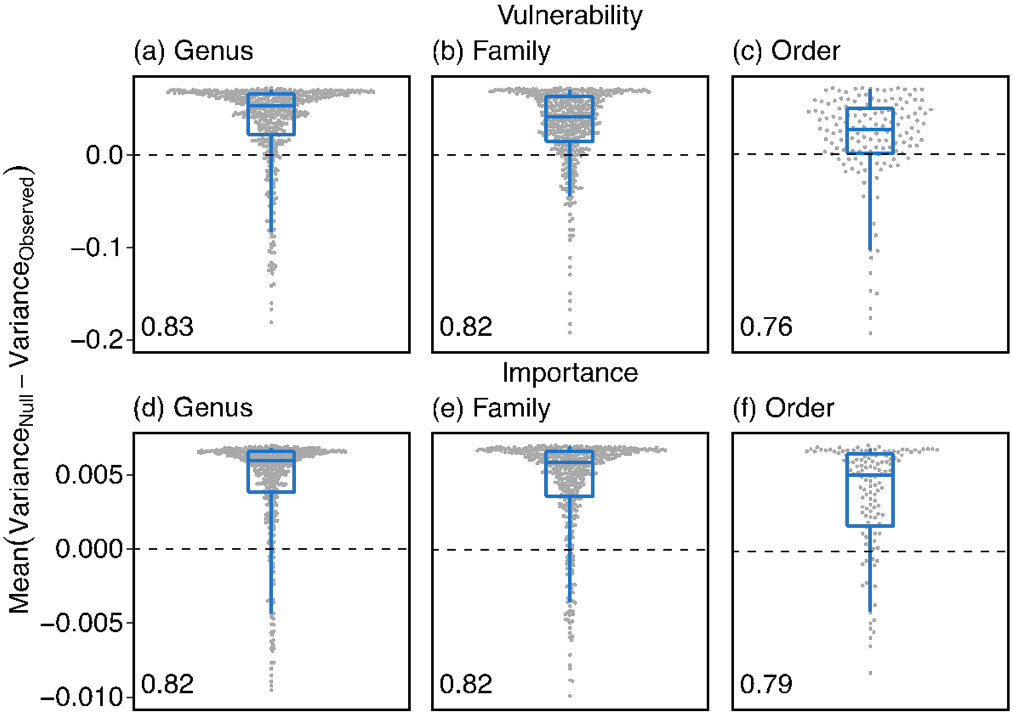


**Figure S2:** The degree of taxonomic consistency for each interaction at each taxonomic level, for both vulnerability (likelihood of a link being lost) and importance (contribution of a link to a network's structural stability). Taxonomic consistency is the tendency for properties of an interaction to be more similar across occurrences than expected by chance. Points represent individual interactions. Boxplots represent 5%, 25%, 50%, 75% and 95% quantiles of the same data, moving from the bottom whisker to the top whisker. Number in bottom left of each panel is the proportion of interactions which exhibited positive consistency ($Variance_{Observed} < Variance_{Null}$). For visualisation, a small number of points with low values were removed. The percentage of points with values lower than the y-axis minimum are as follows for each panel: (a) 1.5%, (b) 1.1%, (d) 7.2%, (e) 6%, (f) 5.3%.
